# Universal scaling of maximum speed with body mass - Why the largest animals are not the fastest

**DOI:** 10.1101/095018

**Authors:** Myriam R. Hirt, Walter Jetz, Bjöern C. Rall, Ulrich Brose

## Abstract

Speed is the fundamental constraint on animal movement, yet there is no general consensus on the determinants of maximum speed itself. Here, we provide a universal scaling model of maximum speed with body mass, which holds across locomotion modes, ecosystem types and taxonomic groups. In contrast to traditional power-law scaling, we predict a hump-shaped relationship due to a finite acceleration time for animals. This model is strongly supported by extensive empirical data (470 species with body masses ranging from 5.7×10^−8^ to 108,000 kg) from terrestrial as well as aquatic ecosystems. Our approach offers a novel concept of what determines the upper limit of animal movement, thus enabling a better understanding of realized movement patterns in nature and their multifold ecological consequences.

---

The movement of animals and its consequences for ecosystem functioning have long fascinated humans and triggered enormous research^1,2^. Nevertheless, a generalized understanding of what determines variation in movement across species and environments is still lacking. This is particularly important as movement is one of the most fundamental processes of life: the individual survival of mobile organisms depends on their ability to reach resources and mating partners, escape predators, and switch between habitat patches or breeding and wintering grounds. Moreover, by creating and sustaining individual home ranges^3^ and meta-communities^4^, movement also profoundly affects the ability of animals to cope with land-use and climate changes^5^. Additionally, movement determines encounter rates and thus the strength of species interactions^6^, which is an important factor influencing ecosystem stability^7^. Thus, a generalized and predictive understanding of animal movement is crucial.

Maximum speed is the fundamental constraint of movement. The realized movement depends on ecological factors such as landscape structure, habitat quality, or sociality, but the range within which this realized movement occurs meets its upper limit at maximum movement speed. Similar to many physiological and ecological parameters, movement speed of animals is often thought to follow a power-law relationship with body mass^8–10^. However, scientists have always struggled with the fact that in running animals the largest are not the fastest. In nature, the fastest animals such as cheetahs or marlins are of intermediate size indicating that a hump-shaped pattern may be more realistic. There have been numerous attempts to describe this phenomenon^11–15^, but a universal mechanistic model explaining this relationship is still lacking. Here, we fill this void by a novel maximum speed model based on the concept that animals are limited in their time for maximum acceleration due to restrictions on the quickly available energy. Consequently, acceleration time becomes the critical factor determining the maximum speed of animals. In the following, we first derive the maximum-speed model (in equations that are illustrated in the conceptional Fig. 1) and, subsequently, test the model predictions employing a global data base and eventually illustrate its applications to advance a more general understanding of animal movement.

**Figure 1.**
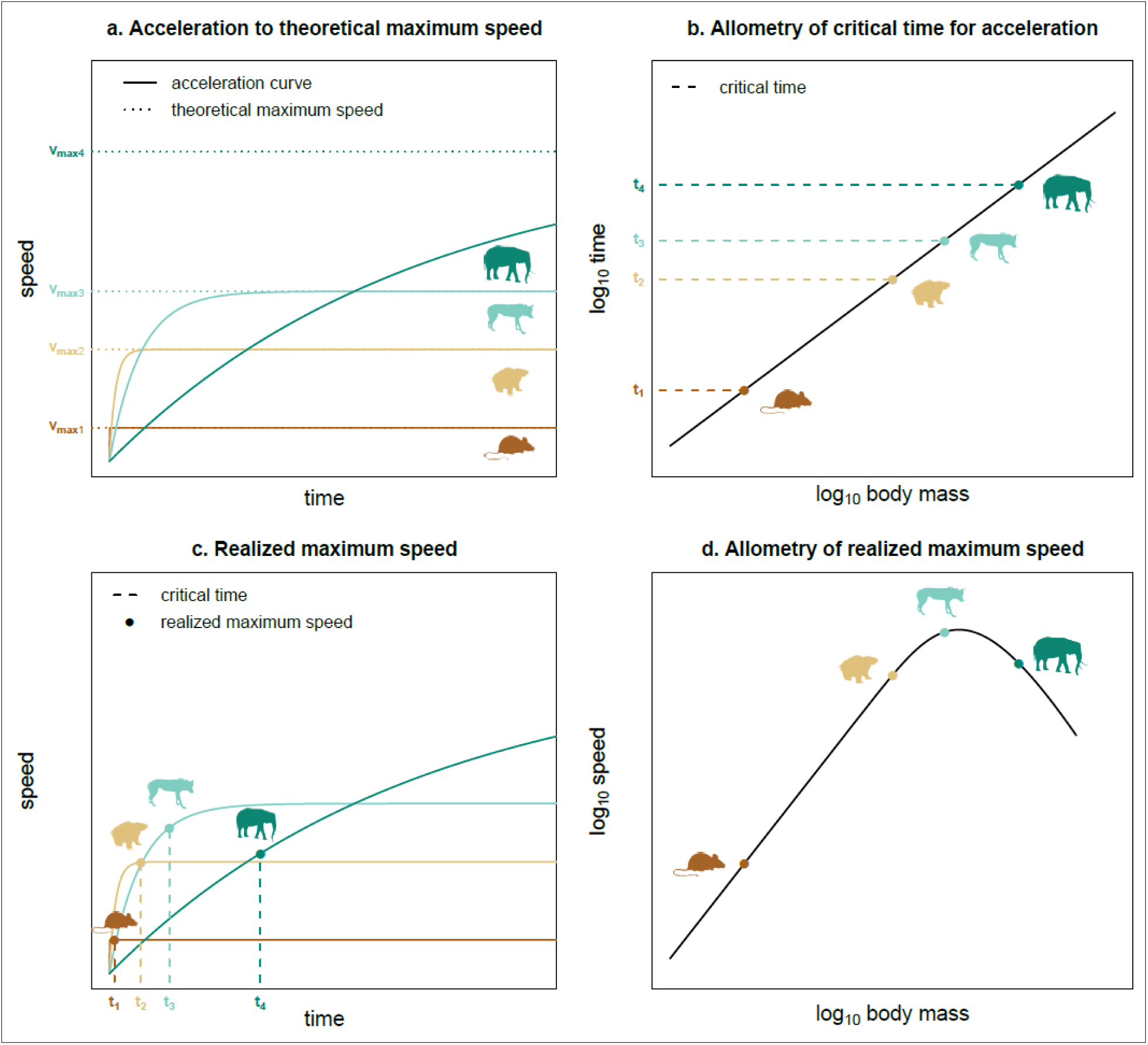
Concept of time- and mass-dependent realized maximum speed of animals. Acceleration of animals follows a saturation curve (solid lines) approaching the theoretical maximum speed (dotted lines) depending on body mass (blue color code) **(a)**. The time available for acceleration increases with body mass following a power law **(b)**. This critical time determines the realized maximum speed **(c)** yielding a hump-shaped increase of speed with body mass **(d)**.

Consistent with prior models, we start with a power-law scaling of theoretical maximum speed v_max(theor)_ of animals with body mass *M*:

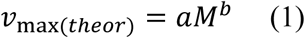

During acceleration, the speed of an animal over time *t* saturates (Fig 1a, solid lines) approaching v_max(theor)_ (Fig 1a, dotted lines):

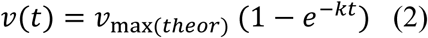

The acceleration constant *k* describes how fast an animal reaches v_max(theor)_. Based on the Newtonian principle *F*=*M* \**k*, acceleration *k* should scale relative to the ratio between maximum force, *F*, and body mass, *M*: *k* ~ *F*/*M*. Knowing that maximum muscle force is roughly proportional to body mass as: *F* ~ *M^d^*, this yields a general power law scaling of *k* with body mass *M*:

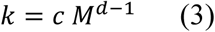

with constants *c* and *d*. As the allometric exponent *d* of the muscle force falls within the range 0.75 to 0.94 ^16–18^ the overall exponent (*d*-1) should be negative, implying that larger animals need more time to accelerate to the same speed than smaller ones (conceptional Fig 1a, color code exemplifies four animals of different size). Note that this general scaling relationship also allows for the special cases of a constant acceleration across species or a linear relationship with body mass.

While prolonged high speeds are related to the maximum aerobic metabolism, maximum burst speeds are linked to anaerobic capacity^19,20^. For maximum aerobic speed, so-called slow twitch fibers are needed, which are highly efficient at using oxygen for generating adenosine triphosphate (ATP) to fuel muscle contractions. Thus, they produce energy more slowly but for a long period of time before they fatigue and allow for continuous, extended muscle contractions. In contrast, maximum anaerobic speed is fueled by a special type of so-called fast twitch fibers, which use ATP from the ATP storage of the fiber until it is depleted. Thus, they produce energy more quickly but also fatigue very fast and only allow for short bursts of speed. Consequently, the critical time available for maximum acceleration τ is limited by the amount of fast twitch fibers and their energy storage capacity. This storage capacity is correlated with the amount of muscle tissue mass, which is directly linked to body mass. Thus, similar to the muscle tissue mass, τ should follow a power law:

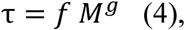

where the allometric exponent *g* should fall in the range 0.76 to 1.27 documented for the allometric scaling of muscle tissue mass ^21–24^ This power-law implies that larger animals should have more time for acceleration (dashed red lines in conceptional Fig 1b and c). However, the power law relationship of the critical time τ in our model allows for a negative or positive scaling of energy availability with body mass as well as the lack of a relationship (constant energy availability across body masses (*f* = 0)). While we included power-law relationships of k and τ (equations 3 and 4) in our model, these scaling assumptions are not strictly necessary. Instead, our only critical assumptions are that acceleration over time follows a saturation curve (equation (1)) and that the time available for acceleration is limited.

Within the critical time τ, after which energy available for acceleration is depleted, the animal reaches its realized maximum speed v_max_ (points in Fig 1c), which may be lower than the theoretical maximum speed (Fig 1a, dotted lines). Combining equations (1) – (4) with t = τ yields 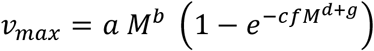 which simplifies to

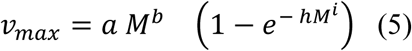

where 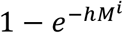 is the limiting factor that determines the realized maximum speed depending on the critical time and the body mass. This equation predicts a hump-shaped relationship between realized maximum speed and body mass (conceptional Fig 1d). Based on the allometric power-law exponents of muscle forces (0.75≤*d*≤0.94) and muscle mass (0.76≤*g*≤1.27), we expect that the exponent *i* (*i*=*d*-1+*g*) should fall in the range between 0.51 and 1.21.

The limiting term 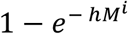 represents the fraction of the theoretical maximum speed that is realized and is defined on the interval] 0;1[. For low body masses, this term is close to 1 and the realized maximum speed approximates the theoretical maximum speed (black and green dots in Fig. 1c). With increasing body masses, this term decreases and reduces the realized maximum speed (blue and yellow dots in Fig 1c). Put simply, small to intermediately sized animals accelerate quickly and have enough time to reach their theoretical maximum speed whereas large animals are limited in acceleration time and run out of readily mobilizable energy before being able to reach their theoretically possible maximum speed. Therefore, they have a lower realized maximum speed than predicted by a power-law scaling relationship.

To test the model predictions (Fig. 1d), we compiled literature data on maximum speeds of running, flying and swimming animals including not only mammals, fish and bird species but also reptiles, mollusks and arthropods. Body masses of these species range from 5.7×10^−8^ to 108,000 kg. Statistical comparison amongst multiple models (see Methods) shows that the time-dependent maximum speed model is the most adequate (see Supplementary Table 3). Our model (Fig. 2, parameter values in Supplementary Table 4) shows that the initial power-law increase of speed with body mass is similar for running and flying animals (*b* = 0.24 and 0.27, respectively). However, flying animals are nearly six times faster than running ones (*a* = 144 and 26, respectively). For swimming animals, the power-law increase in speed is steeper than expected (*b* = 0.36, Fig 2a). This is due to the fact that in contrast to air (in which both flying and running animals move), water is 800 times denser and 60 times more viscous^25^. Small aquatic animals are slower than running animals of the same body size while larger species approach a similar speed as their running equivalents. This implies that in water, size brings a greater benefit in gaining speed. The second exponent is higher for flying animals (*i* = 0.72) than for running (*i* = 0.6) and swimming ones (*i* = 0.56), which fits into the expected range (0.51≤*i*≤ 1.21). Future research will need to disentangle the relative importance of anaerobic and musculoskeletal constraints on movement speed by measuring muscle force, muscle mass, body mass and maximum acceleration for the same species to narrow down this large range of possible exponents. Furthermore, this may allow to address the systematic differences in the exponent *i* between the locomotion modes as well as potential morphological side effects (e.g. quadrupedal vs. bipedal running or soaring vs. flapping flight).

**Figure 2.**
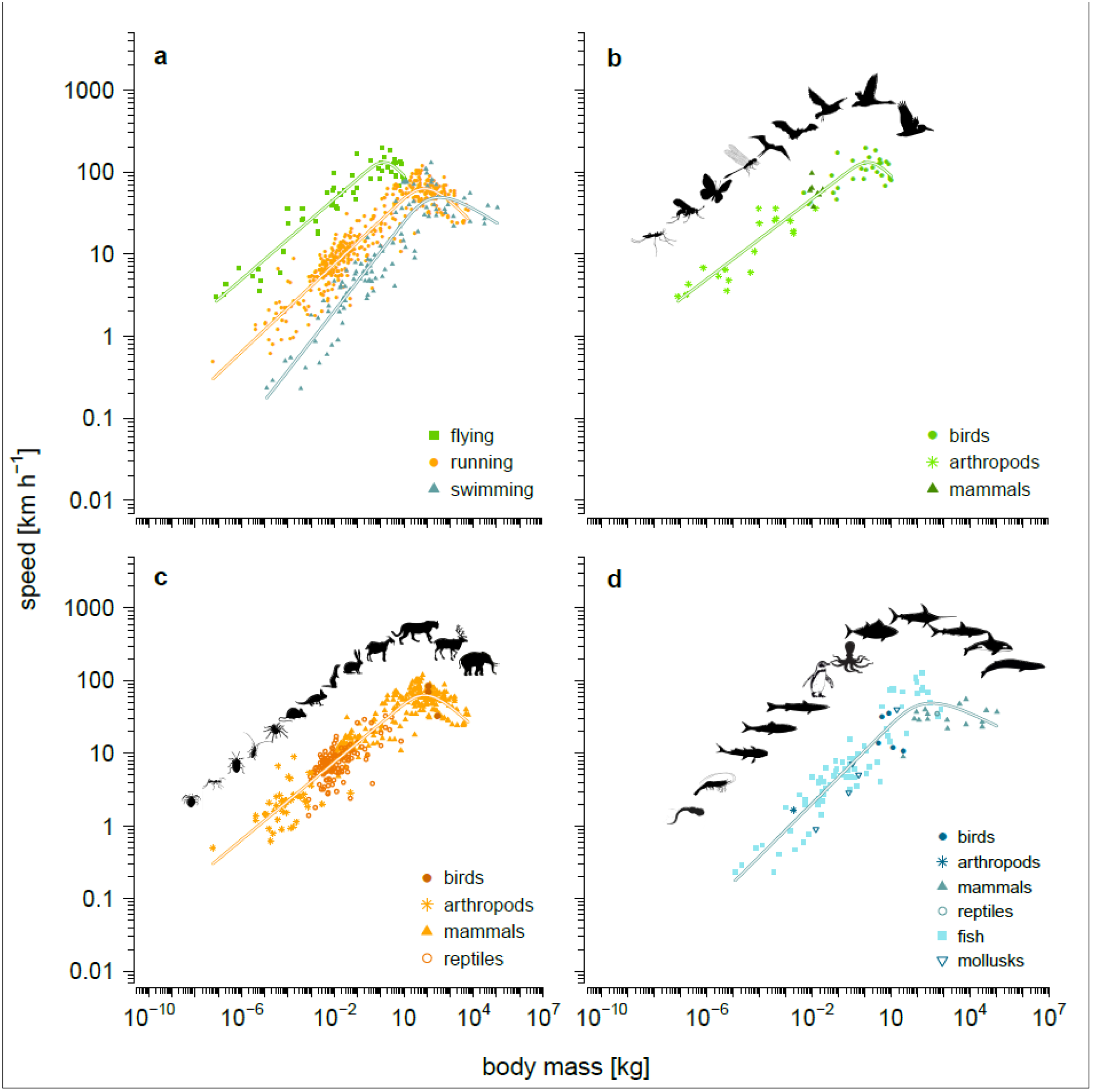
Empirical data and time-dependent model fit on the allometric scaling of maximum speed. **(a)** Scaling for the different locomotion modes (flying, running, swimming) in comparison. Taxonomic differences are illustrated separately for **(b)** flying (n = 55), **(c)** running (n = 453) and **(d)** swimming (n = 109) animals. Overall model fit: R^2^ = 0.893. The residual variation does not exhibit a signature of taxonomy (only a weak effect of thermoregulation, see Methods).

While the model provides strikingly strong fits with observations (R^2^ = 0.893), obvious unexplained variation remains. This might partially be explained by the fact that our data probably include not only maximum *anaerobic* speeds but also some slightly slower maximum *aerobic* speeds. Moreover, we assessed the robustness of our model by exploring this residual variation with respect to taxonomy (arthropods, birds, fish, mammals, mollusks, and reptiles), primary diet (carnivore, herbivore, omnivore), thermoregulation (ectotherm, endotherm) and locomotion mode (flying, running, swimming). As taxonomy and thermoregulation are highly correlated, we made a first model without taxonomy and a second model without thermoregulation and compared them by their BIC values (see Methods for details). According to this, the model including thermoregulation instead of taxonomy is the most adequate (∆BIC = 27.37). In this model, the differences between the diet types were not significant. In contrast, combinations of locomotion mode with thermoregulation exhibited significant differences (Fig. 3). In flying and running animals, endotherms generally tend to be faster than ectotherms (Fig. 3a and b). Metabolic constraints may enable endotherms to have higher activity levels compared to ectotherms at the low to intermediate temperatures most commonly encountered in nature^29^. This pattern is reversed in aquatic systems where endotherms (mammals and penguins) are significantly slower than ectotherms (mainly fish, Fig. 3c). We assume that this is due to the transition from a terrestrial to an aquatic lifestyle aquatic endotherms underwent. Semi-aquatic endotherms are adapted to movement in two different media, which reduces swimming efficiency in comparison to wholly marine mammals: they have 2.4×10^5^ times higher costs of transport^30^. But also in marine mammals, costs of transport are considerably higher than in fish of similar size because they have higher energy expenditures for maintaining their body temperature^30^. Thus, the effect of thermoregulation on the allometric scaling of maximum speed depends on the locomotion mode and the medium. Overall, this significant effect of thermoregulation explained only ~ 4% of the residual variation suggesting that the vast majority of the variation in speed across locomotion modes, ecosystem types and taxonomic groups is well explained by our maximum speed model.

**Figure 3.**
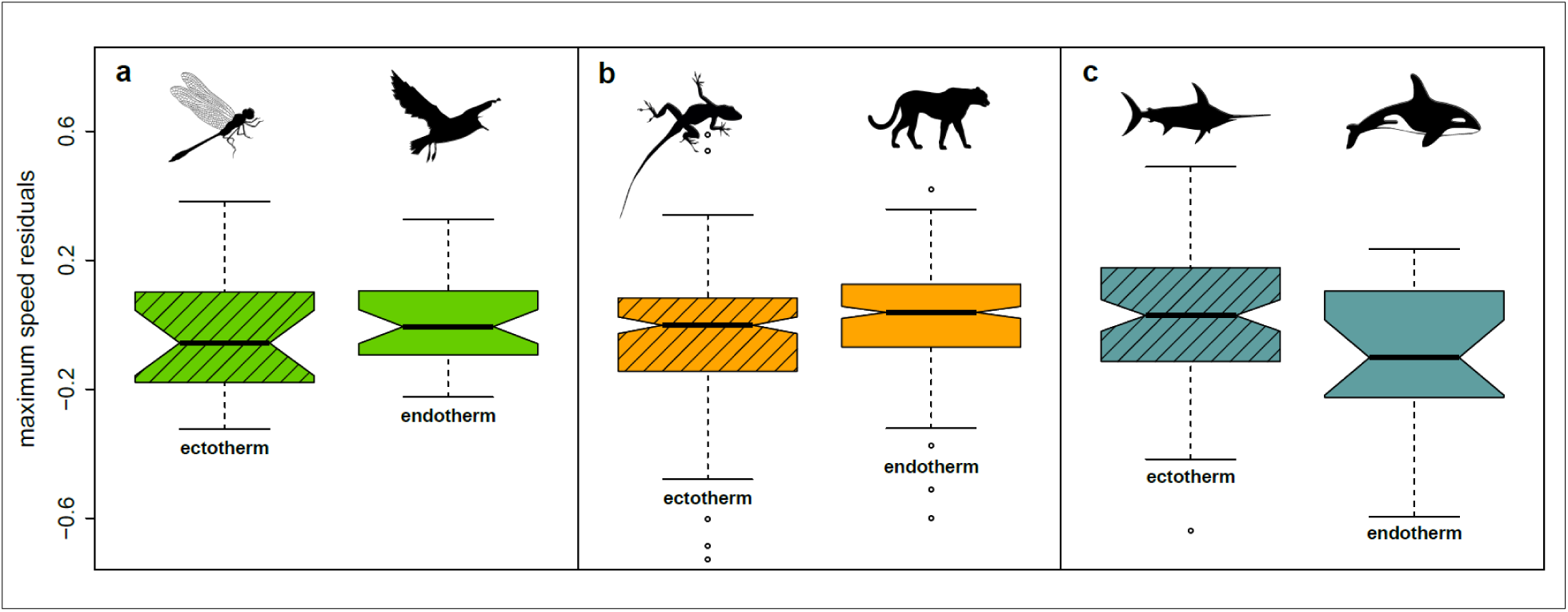
Effect of thermoregulation on the maximum speed of animals (residuals of the relationship in Fig. 2). In flying **(a)** and running **(b)** animals, endotherms are generally faster than ectotherms. In swimming animals **(c)** this effect is reversed with ectotherms being generally faster than endotherms. Box plots show medians (horizontal line), an approximation of 95% confidence intervals suitable for comparing two medians (notches), 25th and 75th percentiles (boxes), the most extreme values within 1.5 times the length of the box away from the box (whiskers), and outliers (dots).

Our findings help solve one of the most challenging questions in movement ecology over the last decades: why are the largest animals not the fastest? Many studies tried to answer this based on morphology, locomotion energetics and biomechanics^10–14^. However, the body-size related hump-shaped pattern observed in empirical data on animal movement speed remained unresolved. To account for this, some studies used polynomials but still lacked a mechanistic understanding^22–24^. Others suggested a threshold beyond which animals run slower than predicted by a power-law relationship due to biomechanical constraints^13^. Therefore, they propose different speed scaling trends depending on the body-mass range^11,12^. Instead of applying different laws to small and large animals, we provide the first universal mechanistic model explaining the hump-shaped relationship between maximum speed and body mass. Our speed predictions are thereby only derived from two major species traits: body mass and locomotion mode, which explain almost 90% (R^2^ = 0.893) of the variation in maximum speed. This general approach allows a species-level prediction of speed which is crucial for understanding movement patterns, species interactions and animal space use. However, our model not only allows prediction of the speed of extant but also that of extinct species. For example, paleontologists have long debated potential running speeds of large birds^31^ and dinosaurs^32,33^ roaming past ecosystems. The benchmark of speed predictions is set by detailed morphological models^32,33^. Interestingly, our maximum speed model yields similar predictions by only accounting for body mass and locomotion mode (Table 1). For instance, in contrast to a power-law model, the morphological and the time-dependent model predict lower speeds of *Tyrannosaurus* compared to the much smaller *Velociraptor*. This is consistent with theories claiming that *Tyrannosaurus* was very likely a slow runner^34^. A simple power-law model only yields reasonable results for lower body masses (e.g. flightless birds) while predictions for large species such as giant quadrupedal dinosaurs are unrealistically high. In contrast, our time-dependent model makes adequate predictions for small as well as large species (almost 80% of the morphological speed predictions are within the confidence intervals of our model predictions, Table 1).

**Table 1.**
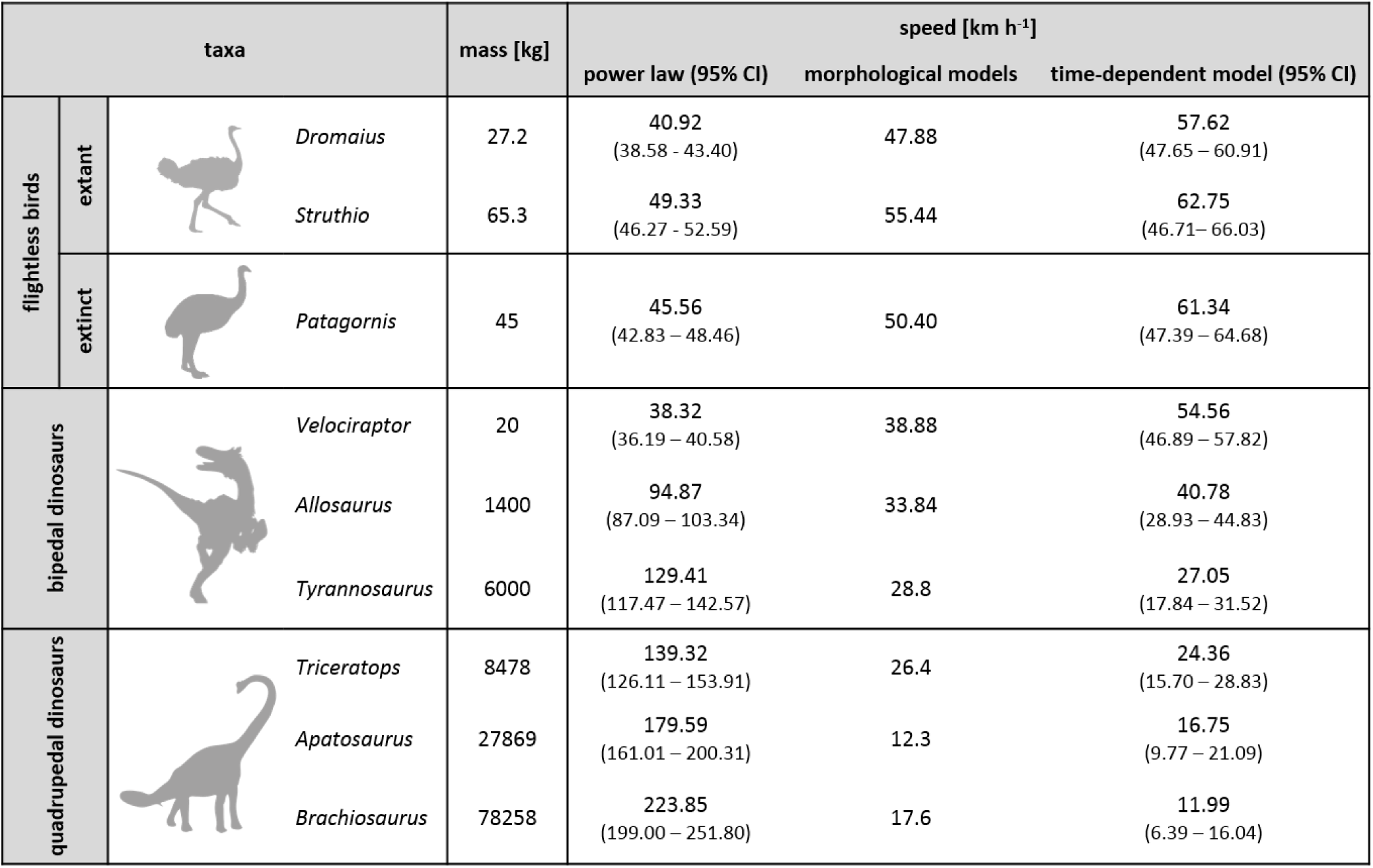
**Maximum speed predictions** of extant and extinct flightless birds and bipedal and quadrupedal dinosaurs. Model predictions of a simple power law, morphological models and our time-dependent maximum speed model are compared (references in Supplementary Table 5). 95% confidence intervals (CI) are given for the power law and time-dependent model.

| taxa |  |  | mass [kg] | speed [km h<sup>-1</sup>] |  |  |
| --- | --- | --- | --- | --- | --- | --- |
|  |  |  |  | power law (95% CI) | morphological models | time-dependent model (95% CI) |
| flightless birds | extant | *Dromaius* | 27.2 | 40.92<br>(38.58 - 43.40) | 47.88 | 57.62<br>(47.65 - 60.91) |
|  |  | *Struthio* | 65.3 | 49.33<br>(46.27 - 52.59) | 55.44 | 62.75<br>(46.71 - 66.03) |
|  | extinct | *Patagornis* | 45 | 45.56<br>(42.83 - 48.46) | 50.40 | 61.34<br>(47.39 - 64.68) |
| bipedal dinosaurs |  | *Velociraptor* | 20 | 38.32<br>(36.19 - 40.58) | 38.88 | 54.56<br>(46.89 - 57.82) |
|  |  | *Allosaurus* | 1400 | 94.87<br>(87.09 - 103.34) | 33.84 | 40.78<br>(28.93 - 44.83) |
|  |  | *Tyrannosaurus* | 6000 | 129.41<br>(117.47 - 142.57) | 28.8 | 27.05<br>(17.84 - 31.52) |
| quadrupedal dinosaurs |  | *Triceratops* | 8478 | 139.32<br>(126.11 - 153.91) | 26.4 | 24.36<br>(15.70 - 28.83) |
|  |  | *Apatosaurus* | 27869 | 179.59<br>(161.01 - 200.31) | 12.3 | 16.75<br>(9.77 - 21.09) |
|  |  | *Brachiosaurus* | 78258 | 223.85<br>(199.00 - 251.80) | 17.6 | 11.99<br>(6.39 - 16.04) |

Our model also allows drawing inferences about evolutionary and ecological processes by analyzing the deviations of empirically measured speeds from the model predictions. Higher maximum speeds than predicted indicate evolutionary pressure on optimizing speed capacities that could for instance arise from co-evolution of pursuit predators and their prey. In ecological research, our maximum-speed model provides a mechanistic understanding of the upper limit to animal movement patterns during migration, dispersal or bridging habitat patches. The travelling speed characterizing these movements is the fraction of maximum speed that can be maintained over longer periods of time. The integration of our model as a species-specific scale (“what is physiologically possible”) with research on how this fraction is modified by species traits and environmental parameter such as landscape structure, resource availability and temperature (“what is ecologically realized in nature”) can help provide a mechanistic understanding unifying physiological and ecological constraints on animal movement. In addition to generalizing our understanding across species traits and current landscape characteristics, this integrated approach will facilitate the prediction of how species-specific movement and subsequently home ranges and meta-communities may respond to the ongoing landscape fragmentation and environmental change. Our approach may act as a simple and powerful tool for predicting the natural boundaries of animal movement and help gain a more unified understanding of the currently assessed movement data across taxa and ecosystems^1,2^.

## Methods

### Data

We searched for published literature providing data on the maximum speeds of running, flying and swimming animals by employing the search terms “maximum speed”, “escape speed” and “sprint speed”. From this list, we excluded publications on (1) vertical speeds (mainly published for birds) to avoid side-effects of gravitational acceleration that are not included in our model or (2) the maxima of normal speeds (including also dispersal and migration). This resulted in a data set containing 617 data points for 470 species (see Supplementary Table 1 for an overview). Our data include laboratory and field studies as well as meta-studies (which are mainly field studies but may also include a minor amount of laboratory studies). For some data points, the study type could not be ascertained and they were marked as “unclear”. For an overview of the study type of our data see Supplementary Table 2.

### Data availability

The full data set is available on the iDiv data portal.

### Model fitting

We fitted several models to these data: (1) the time-dependent maximum speed model (equation (5)), (2) three polynomial models (a. simple polynomial model without cofactor, b. polynomial model with taxon as cofactor but without interaction term and c. polynomial model with taxon as cofactor with interaction term) and (3) three power law models (a. simple power law without cofactor, b. power law with taxon as cofactor but without interaction term and c. power law with taxon as cofactor with interaction term). For swimming animals, we excluded reptiles and arthropods from the statistical analyses as they only contained one data point each (see Supplementary Table 1). The polynomial and power law models were fitted by the lm function and the time-dependent model by the nls function in R 3.2.3^<u>35</u>^. The quality of the fits was compared according to the Bayesian information criterion (BIC) that combines the maximized value of the likelihood function with a penalty term for the number of parameters in the model. The model with the lowest BIC is preferred, which demonstrates that the time-dependent maximum speed model developed in the main text was most adequate in all cases (see Supplementary Table 3). For flying animals, the simple polynomial model performed second best, whereas for running animals the polynomial model with taxon as cofactor with interaction term and for swimming animals the power-law model with taxon as cofactor with interaction term were second best (see Supplementary Table 3). Overall, the lower BIC values indicate that the time dependent maximum speed model provides a fit to the data that is substantially superior over power-law relationships, models with taxonomy as cofactor or (non-mechanistic but also hump-shaped) polynomials. The fitted parameter values of the time-dependent maximum speed model for flying, running and swimming animals are given in Supplementary Table 4.

### Residual variation analysis

We analyzed the residuals of the time-dependent maximum-speed model (Fig. 2 of the main text) with respect to taxonomy (arthropods, birds, fish, mammals, mollusks, and reptiles), primary diet type (carnivore, herbivore, omnivore), locomotion mode (flying, running, swimming) and thermoregulation (ectotherm, endotherm) using linear models. As taxonomy and thermoregulation are highly correlated, we made a first model without taxonomy and a second model without thermoregulation:

Model 1: residuals ~ (thermoregulation + diet type) * locomotion mode
Model 2: residuals ~ (taxonomy + diet type) * locomotion mode

We compared both models via BIC and carried out a further mixed effects model analysis on the superior model. This model included the study type as a random factor influencing the intercept, which ensures that differences among study types do not drive our statistical results. We acknowledge that the direct inclusion of multiple covariates in the model-fitting process would be preferable over residual analysis to avoid biased parameter estimates^36^. However, this was impeded by the complexity of fitting the non-linear model with four free parameters (equation (5)), and our main goal was less exact parameter estimation than documenting the main variables affecting the unexplained variation.

## Acknowledgements

M.R.H, W.J., B.C.R., and U.B. gratefully acknowledge the support of the German Centre for integrative Biodiversity Research (iDiv) Halle-Jena-Leipzig funded by the German Research Foundation (FZT 118).

## Author Contributions

M.R.H. and U.B. developed the model. M.R.H. gathered the data. M.R.H. and B.C.R. carried out statistical analyses. W.J. was involved in study concept and data analyses. M.R.H. and U.B. wrote the paper. All authors discussed the results and commented on the manuscript.

